# The association of catechol-O-methyltransferase genetic polymorphism rs4680 in physical activity among adult women

**DOI:** 10.1101/2022.06.13.495863

**Authors:** Lilach Gotlieb, Sigal Ben-Zaken

**Affiliations:** Wingate Academic College, The Genetic and Molecular Biology Laboratory, Netanya, Israel, 4290200

**Keywords:** Physical Activity, Women, *COMT* A/G rs4680

## Abstract

**Background:** The aim of the study was to explore associations between *COMT* A/G rs4680 polymorphisms and physical activity (PA) among healthy middle-aged women. PA is a multifactorial trait in which dopamine plays a pivotal role. The enzyme catecholamine O-methyl transferase (COMT) degrades dopamine in the synaptic area. The *COMT* rs4680 genetic polymorphism results in either *COMT* Met allele or *COMT* Val allele. This functional polymorphism causes differences in enzyme activity with low enzymatic activity (and higher dopamine levels), associated with the Met allele; high enzymatic activity is associated with the Val allele.

**Method:** Ninety healthy women, aged 47±5.5 years from similar demographic backgrounds, participated in the study. PA engagement was assessed by the BAECKA questionnaire of habitual physical activity PA. Genomic DNA was extracted from buccal epithelial cells for *COMT* rs4680 analysis.

**Results:** Despite a similar demographic background of the participants, a large variance was found in all PA indexes. A-allele carriers’ prevalence was the significantly higher (83%) among highly active women compared to its prevalence among moderate (64%) and low (47%) active women.

**Conclusions:** It seems that *COMT* A/G rs4680 A-allele carriers might be associated with a relatively high rate of PA practitioners in general and running in particular

## Introduction

There is a worldwide consensus regarding the physiological and psychological health benefits of physical activity (PA)^1–3^. Yet according to the World Health Organization (WHO), PA engagement among the general population varies greatly, ranging from sedentary to highly active behavior^4– 6^. Women specifically do not meet the recommended weekly physical activity threshold (31.7% are physically inactive) compared to men (23.4% are physically inactive)^1^ – although these number vary greatly across countries^7^.

The complex PA phenomenon is influenced by genetic factor, environmental factors, and their complex interactions. Studies on twins and families estimate a PA heritability that ranges from 20% to 92%, usually converging into 50-60%^8^. Several genetic polymorphisms have been suggested as a source for such PA engagement variability^9^, such as genetic polymorphisms that relate to the neural reward system, which seems to play pivotal role in this range. The neural reward system comprises a number of neural pathways in the brain, which create a sense of pleasure in response to certain stimuli^10^, among other functions. In other words, this system influences behavior by “reinforcing” behaviors that lead to positive rewards, thereby increasing the likelihood of repeating them in the future^11^. A positive response from PA has been found to impact the continued performance of the same behavior, stemming from the desire to receive the positive reward. Thus, feelings of reward or enjoyment are considered significant factors when engaging in PA^12^. People will even increase such activity in search of that reward and sense of pleasure^13^. Therefore, neurotransmitters in the reward system, such as dopamine, serotonin, and norepinephrine, act as modulators of PA behavior and play a role in PA participation^14,15^.

The fundamental role of dopamine in PA makes variants in gene encoding of key proteins in the dopaminergic system natural candidates for explaining PA variability. The enzyme COMT degrades dopamine in the synaptic area. COMT is encoded by the *COMT* gene, settled in the 22q11.2 chromosome. The single nucleotide polymorphism (SNP) rs4680 in the *COMT* gene is formed in the 158^th^ codon of the gene, resulting in either *COMT* Met allele or Val allele^16^. This functional polymorphism causes differences in enzyme activity that are threefold or even fourfold, with low enzymatic activity being associated with the Met allele, and high enzymatic activity being associated with the Val allele^17^. Moreover, the slow degradation of dopamine by the *COMT*-Met allele results in higher dopamine levels^17^. Finally, Cerebral dopaminergic function regulates several factors relating to PA, including cognitive control^18^, physical function^19^, depressive moods^20^, and motivation and reward responses^21,22^.

Some observational studies regarding associations between *COMT* rs4680 polymorphism and self-reported PA, however, were inconclusive in their findings^16,23^, which could be the result of a large age range among participants and studies that included both males and females combined. The aim of the current study, therefore, was to explore associations between *COMT* A/G rs4680 polymorphisms and PA among healthy middle-aged women, from similar socio demographic backgrounds. We hypothesized that *COMT* A/G rs4680 A-allele, which is associated with high level of prefrontal dopamine, will be more prevalent among active women – particularly among women whose main PA is running.

## Materials and Methods

### Participants

Ninety healthy women, aged 47±5.5 years participated in the study. These women in responded to ads published in various media, in addition to the “friend brings friend” method. All participants lived in urban areas in Israel, were mainly married (82%) with children (92%), from high socio-demographic statuses (80% had an academic education; 66% had an above-average income). Of the 90 women, 79 indicated that they engaged in some form of PA (88% were ‘active’), of whom 46 reported their main activity to be running (58%, ‘active-runners’ [AR]); 33 indicated being active in other sport activities, such as walking, Pilates or yoga (42%, ‘active non-runners’ [ANR]). The Active participants were divided into three groups, according to their weekly exercise frequency (WEF): (1) Low (up to two hours per week); (2) moderate (two-three hours per week); and (3) high (more than three hours per week).

The study was approved by the Institutional Review Board of the Hillel Yaffe Medical Center, Hadera, Israel, Helsinki certificate number HYMC 111-16. A signed written informed consent was obtained from each participant.

### PA assessment

The Baecke Physical Activity Questionnaire (BPAQ) was used for assessing the participants’ PA. This questionnaire was designed to quantify work, sports, and non-sports leisure activities, and was found to be valid and reliable^24–26^. The BPAQ has a total of 16 questions and consists of three parts: Work-related PA section included questions about the physical demands of work, the Sport-related PA section included questions about the frequency and intensity of participating in sports. And Leisure-time PA section included questions about everyday PA outside of work and sports. The answers were based on a 5-point Likert-like scale and were used to calculate an index for each section: The Occupation Activity Index (OAI), Sport Activity Index (SAI), and Leisure Time Activity Index (LAI). In addition, a running section was added to the questionnaire, dealing with women’s running habits during leisure time. A Running index (RI) was calculated.

### Genotyping

Genomic DNA was extracted from buccal epithelial cells collected using a cytological brush. DNA was extracted using a commercial kit (Qiagen Buccal Cell and DNAeasy Kit), as per the manufacturer’s instructions. The Assayb y-Design service (www.appliedbio-systems.com) was used to set up a Taqman allelic discrimination assay for the *COMT* A/G SNP rs4680.

The following primer sequences were applied: *forward:* TGAGTCCCTGAACCAGCAAAG, and *reverse:* GACGTGCCCACCTGTGAT. The Polymerase Chain Reaction (PCR) mixture included 5 ng genomic DNA, 0.125 μl TaqMan assay (40*, ABI), 2.5 μl Master mix (ABI), and 2.375 μl water. PCR was performed in 384 well PCR plates in an ABI 9700 PCR system (Applied Biosystems Inc., Foster City, CA, USA), and consisted of initial denaturation for 10 min at 95°C, 40 cycles with denaturation of 15 s at 92°C, and annealing and extension for 60 s at 60°C. Results were analyzed by the ABI Taqman 7900HT using the sequence detection system 2.22 software (Applied Biosystems Inc.).

### Demographic and Anthropometric measurements

All participants answered a demographic questionnaire and anthropometric measurements were taken: waist circumference (WC) Hip circumference (HC). The measurements were done using a soft, non-stretchable measuring tape. The circumferences were recorded in centimeters. The measurement was made according to a protocol of the World Health Organization^27^.

### Data analysis

The SPSS statistical package, version 20.0, was used to perform all statistical analysis (SPSS, Chicago, IL, USA). A χ2-test was used to confirm that the observed genotype prevalence was within the Hardy-Weinberg equilibrium. A χ2-test was used to compare alleles and genotypes prevalence between: ‘active’ and ‘non-active’, between different groups of WEF, and between AR and the ANR groups. One-way ANOVA analysis was used to compare the PA indexes among genotype carriers. In cases of non-normal variable distribution, a Wilcoxon two-sample test was performed. A two-way ANOVA was performed to compare each anthropometric variable (as dependent values) and the PA indexes, the genotype carriers (as independent values), and the interaction between them. A two-way ANOVA was used to compare the PA indexes and the genotype carriers and the diacritical variable between women who menstruate and women who do not. Statistical significance was adjusted for multiple testing.

## Results

### PA variability

Our findings indicate a large variability in PA performed during the daytime, with the PA index ranging from 3.5AU to 14.2AU (mean 2.02, SD=2.45). Of the three PA indexes, the greatest variability was found in the SAI (SD=1.56 compared to SD=0.76 in the other indexes). The PA index distribution is presented in Figure 1. The SAI index, which presented the largest distribution, revealed a one-end tail, with more participants engaging in none-to-low levels of sports activities. The LAI exhibited normal distribution.

**Figure 1.**
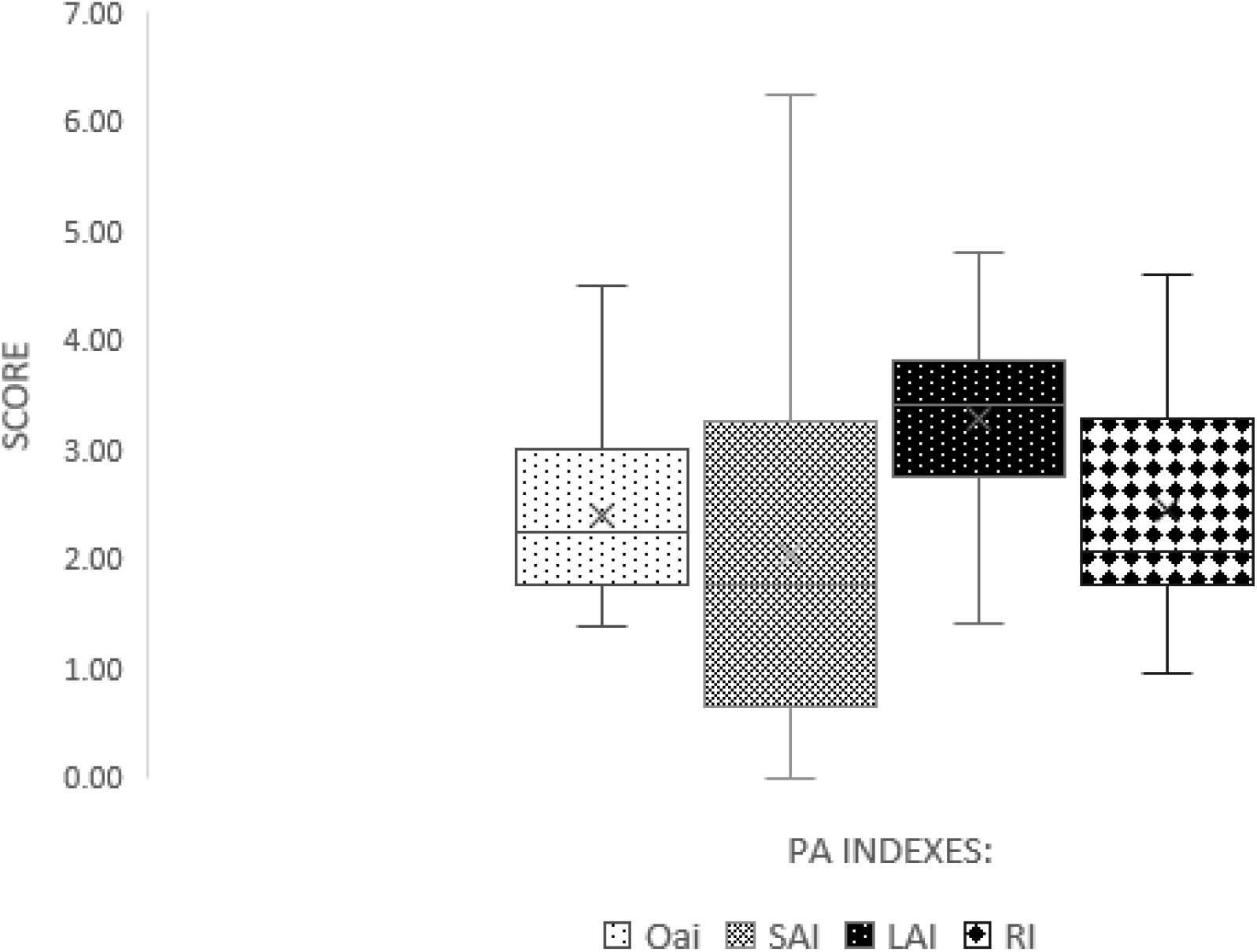
PA index distribution. The SAI has the largest distribution: largest range and largest interquartile range (IQR), with a negative asymmetric distribution: the highest spread in the last two quartile. This indicates a very high variance in sports activities. In contrast, distribution of the LAI is normal.

### *COMT* A/G rs4680 prevalence by WEF

Complete data regarding prevalence of genotypes and alleles is presented in Table 1. Significant differences were found in the prevalence of both genotypes (Figure 2A) and alleles (Figure 2B) by WEF. The prevalence of A-allele carriers was highest among women who exercise at high-frequency (83%), compared to its prevalence among women who conduct moderate (64%) and low WEF (47%) (χ^2^_df=2_= 7.23, p = .02 mutant model). A-allele prevalence was higher among women with in high (56%) and moderate (44%) WEF compared to those with low (32%) WEF (χ^2^_df=2_=5.24, p = .06 allele model), as presented in Figure 2.

**Table 1.**
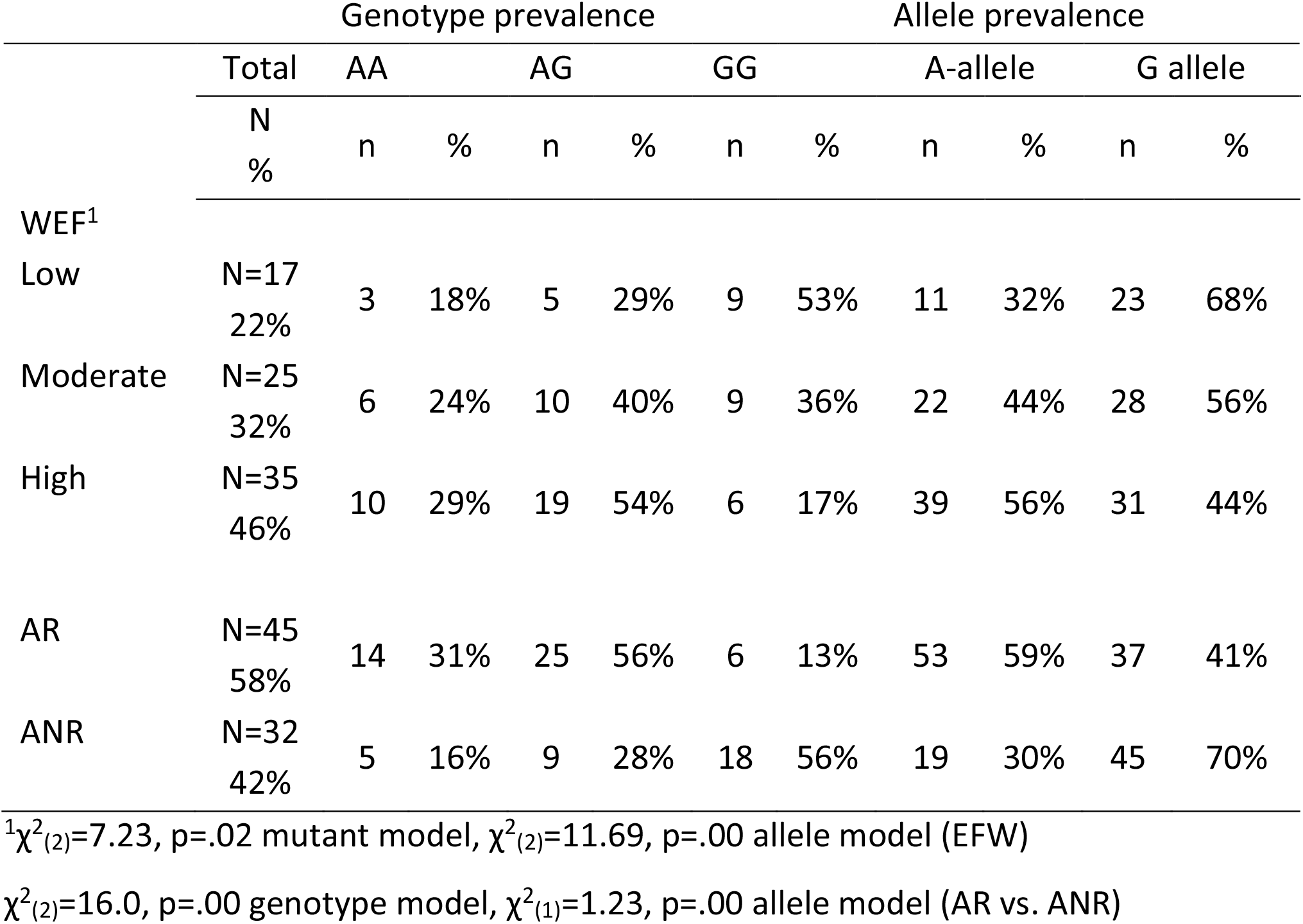
COMT A/G rs4680 genotypes and allele prevalence among Israeli women

**Figure 2.**
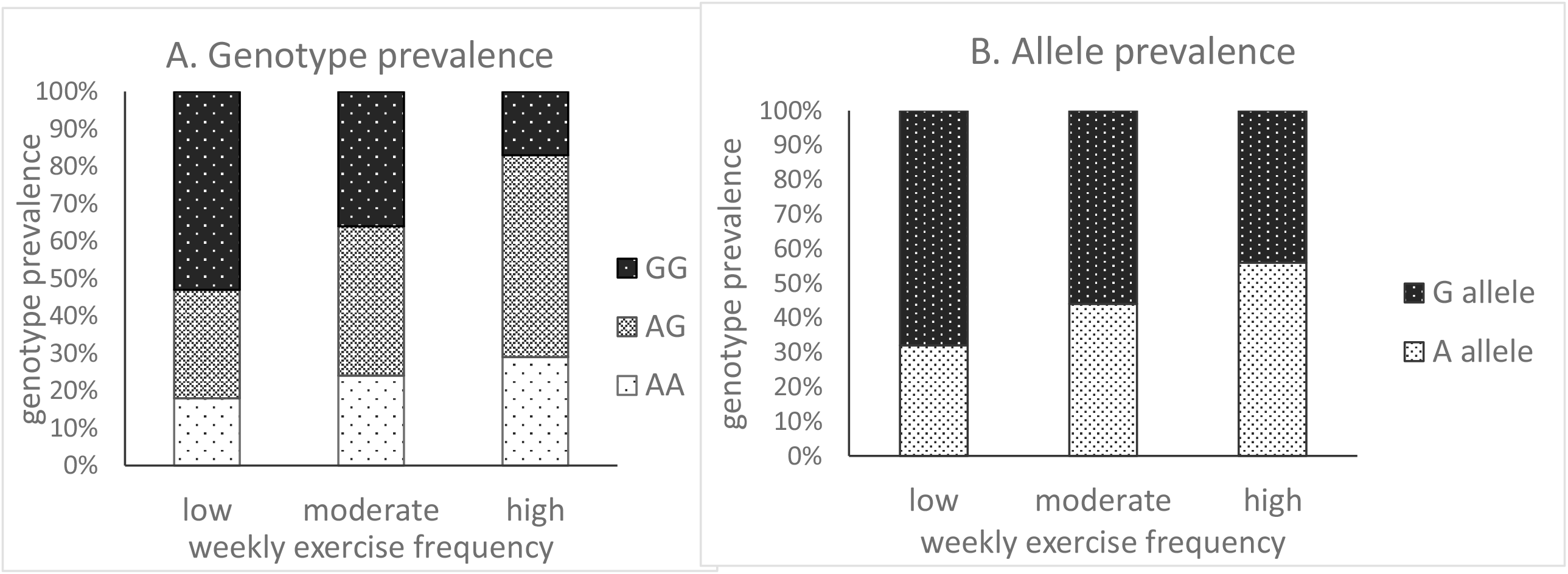
COMT A/G rs4680 genotype (A) and allele (B) prevalence among adult Israeli women by WEF

The prevalence of A-allele carriers was significantly higher among AR (31%-AA, 56%-AG) vr. ANR (16%-AA, 28% AG) (χ^2^_df=2_= 16.0, p = .00 genotype model) as presented in Figure 3. Finally, *COMT* rs4680 prevalence did not differ by age, education, income, Body Mass Index (BMI), WC and HC.

**Figure 3.**
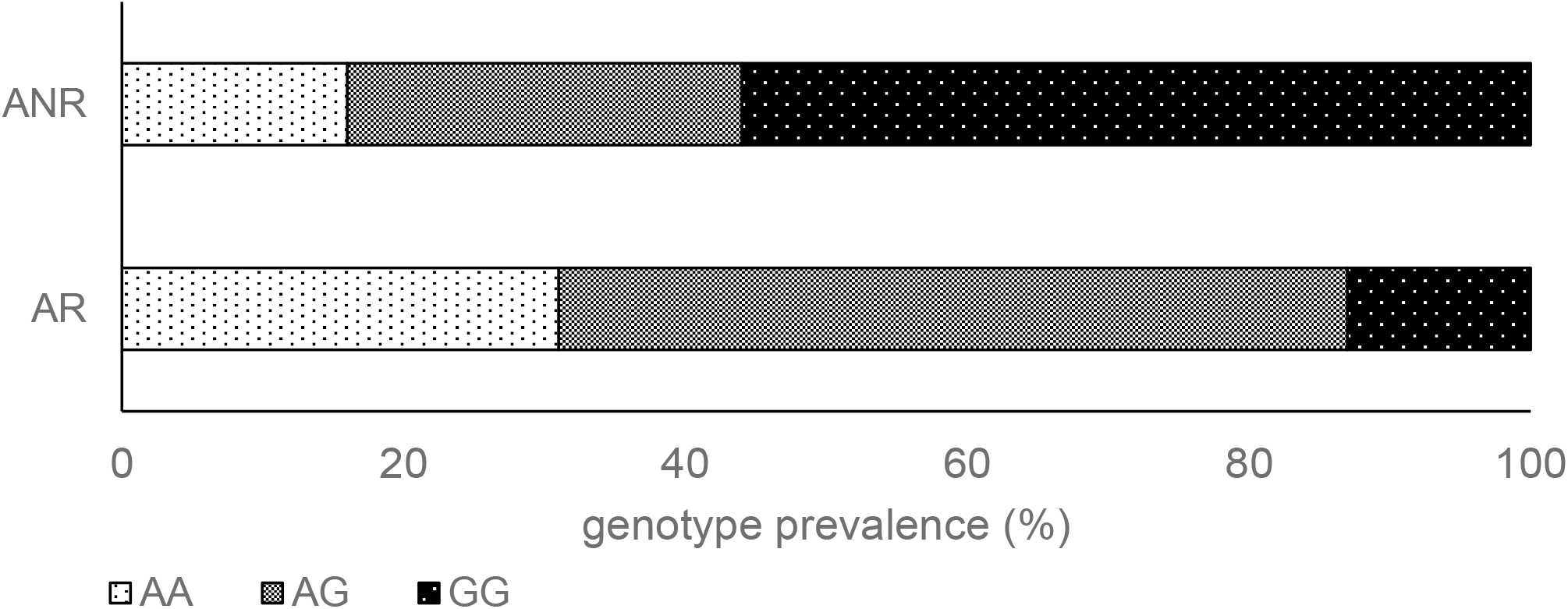
COMT A/G rs4680 genotype prevalence among adult Israeli women divided into AR vs. ANR

### Physical activity indexes according to *COMT* A/G rs4680 polymorphism

Means of the PA indexes divided to *COMT* A/G rs4680 genotypes are presented in Table 2. SAI, LAI and the RI means were significantly higher among A-allele carriers compared to non-carriers. The SAI mean of A-allele carriers was found to be twice as high than the same index for the GG genotype.

**Table 2.** PA indexes: Means and SD divided into COMT A/G rs4680 genotypes and mutant model

| COMT A/G | AA | AG | AA+AG | GG |
| --- | --- | --- | --- | --- |
| rs4680 | n=22 | n=38 | n=60 | n=27 |
| PA Indexes | Mean±SD | Mean±SD | Mean±SD | Mean±SD |
| OAI | 2.16±0.67 | 2.56±0.81 | 2.41±0.78 | 2.35±0.69 |
| SAI | 2.29±1.47<sup>1</sup> | 2.45±1.66 | 2.39±1.58<sup>2</sup> | 1.27±1.20 |
| LAI | 3.29±0.77<sup>3</sup> | 3.48±0.71 | 3.41±0.73<sup>4</sup> | 2.98±0.74 |
|  | n=14 | n=26 | n=40 | n=8 |
| RI | 2.45±0.92<sup>5</sup> | 2.46±1.25 | 2.45±1.13<sup>6</sup> | 1.39±0.99 |

<sup>1</sup> $\chi^2_{df=2}=9.73$ . p=.01 Kruskal-Wallis
<sup>2</sup> U=472. P=.00 Mann-Whitney
<sup>3</sup> $F_{df=2}=3.53$ . p=.03 one way anova
<sup>4</sup> $T_{df=85}=2.48$ . p=.01 Independent t-test
<sup>5</sup> $F_{df=2}=2.59$ . p=.06 one way anova
<sup>6</sup> $T_{df=46}=2.46$ . p=.02 Independent t-test

### WEF and PA indexes by *COMT* A/G rs4680 and menopause status

The participants were divided into two groups: before menopause (BM (77%, n=69)) vs. menopause (ME (23%, n=21)) according to their response to the question “Do you get your period regularly?”. No significant differences were found in *COMT* rs4680 prevalence between the two groups according to the WEF. However, among BM group, 88% (n=22) of the highly active participants and 68% (n=11) of the moderately active participants were A-allele carriers compared to 47% (n=7) of the low active participants (Chi-Sq_df=2_=7.93, p=.01). No difference was found in *COMT* rs 4680 genotype prevalence according to WEF among ME group. In addition, PA indexes (SAI and LAI) were significantly higher among A-allele carriers (mean=2.37, SD=1.67, mean=3.40 SD=0.77 respectively compared to non-carriers (mean=1.08 SD=1.25, mean 2.74 SD=0.68 respectively, only among BM group.

## Discussion

PA engagement results from various inter-related factors, including genetic variability^14,28^, physiological responses^29^, and the social environment^30^. However, even among individuals living in the same environments, considerable individual variability can be seen in PA behavior^31^ – as seen in the current study. The greatest variability was found in sports activity (SAI), while the lowest variability was found in daily life activity (LAI). However, in contrast to daily life activity, sports activities involve conscious choices, decisions, and actions.

Bryan et al., suggests two principal phenotypes relating to PA and sedentary behavior: the relative reinforcing value and the affective response^32^. The core principle behind the relative reinforcing value of PA is the choice between being active or remaining sedentary^33,34^ or in other words, how much work an individual is willing to invest in order to obtain particular active or sedentary behavior^35^. The theoretical concept of the reinforcing value phenotype suggests different neurobiological systems for *wanting* (dopaminergic) versus *liking* (opioid), and as such can provide insights into the biological mechanisms that promote PA^36–38^.

The second phenotype is affective response to exercise, relating to feelings of pleasure or displeasure that people experience while performing PA^39^. The affective response to exercise might result from the interplay of two systems: the cognition system, which involves appraisals, motivation, and goals relating to the activity; and the physiological system, that stems from interoceptive cues that the body experiences in response to an exercise bout^40^.

Multiple studies indicate genetic factors that are associated with PA behavior in general^13,41–43^ and with relative reinforcing value^44^, and affective response to PA^13,45,46^ in particular. Due to the pivotal role of the dopaminergic system in these PA behavior phenotypes, genetic variants related to the dopaminergic system are natural candidates for explaining PA behavioral variability. As dopamine is critical for motor system functioning^47–49^, its synthesis seems to affect people’s will to practice PA^50,51^.

The central function of the dopaminergic system is to control motivation for natural rewards and motor movement^52^. Several animal studies suggest that some aspects of dopaminergic functioning may contribute to the genetic/biological regulation of PA^53–55^. The midbrain dopamine system has been shown to mediate rewards and reinforcements for several behavioral functions. For example, Salamone^56^ suggests that the dopamine system may be intricately involved in both motor control and rewards, with the complexity of this regulation allowing for the modulation of complex motor processes. It has also long been thought that the dopamine system plays a role in the rewarding and/or reinforcement of behavioral functions^57^, and it is suggested that the dopamine system plays a major role specifically in the relative reinforcing value^58^. Thus, dopamine may be an interface between motivation and action^59^, and is a likely candidate for regulating voluntary PA.

Genetic polymorphism that is related to the dopamine neural reward system is a natural candidate for explaining PA variability^16^. Indeed, dopamine related genetic variants have been linked to differences in PA among rodents^60^ and among humans^12^. However, genetic association studies have not been consistently successful in uncovering the causative variants for the initiation and maintenance of voluntary exercise behavior^60,61^. While some studies found a significant association between genetic variants in the dopaminergic system and exercise behavior in humans^62–64^, others have failed to find an association between genes involved in the dopaminergic system and either daily PA or the narrower trait of voluntary exercise^65^. This inconsistency might be attributed to methodological issues, such as PA definition and measurements^66,67^.

The enzyme COMT degrades catecholamines, including dopamine. Two main COMT protein isoforms are known: In most assayed tissues, a soluble cytoplasmic (S-COMT) isoform predominates^68^, while in the brain, a longer membrane-bound form (MB-COMT) is the major species^69^. Dopamine degradation by COMT is especially profound in the prefrontal cortex (PFC)^70,71^. COMT is encoded by the *COMT* gene, which lies on chromosome 22q11. The commonly present G>A polymorphism produces a valine-to-methionine (Val/Met) substitution at codons 108 and 158 of S-COMT and MB-COMT, respectively^72^, that results in a trimodal distribution of COMT activity in human populations^72–74^. The polymorphism is usually referred to as the Val/Met locus, and its reference sequence is rs4680. *COMT* A/G rs4680 polymorphism regulates the amount of dopamine in the prefrontal cortex^16^. *COMT* rs4680 met/met (AA) is associated with higher reward responsiveness compared to other *COMT* rs4680 G allele carriers^22^, probably due to high level of prefrontal dopamine. The association between *COMT* A/G rs4680 polymorphism and PA engagement has been seen in some studies, yet while some studies did not reveal associations between *COMT* rs4680 polymorphism and self-reported PA^16,23^, others reported that *COMT* rs4680 AA Genotype carriers exhibit increased engagement in sport activities in response to intervention programs compared to G-allele carriers^75^.

The main finding of the current study is the significant association between *COMT* rs4680 A/G and PA engagement. *COMT* rs4680 A-allele carriers exhibit significantly higher levels of sport activity and leisure time PA compared to A-allele non-carriers, especially among women whose main sport activity was running. Moreover, more than half of the highly active participants were A-allele carriers, compared to less than of third of the low active participants. Furthermore, it was found that the prevalence of A-Allele carriers among AR was significantly higher compared to their prevalence among ANR. In addition, SAI, LAI, and RI were significantly higher among A-Allele carriers compared to the GG genotype. As such, it seems that A-allele might be associated with increased PA in general and with increased running in particular. It is possible that among female A-allele carriers, there is higher sensitivity to the rewards of PA, especially through running. The sport of running comprises several physical and mental health benefits^76^, and recent years have seen an increase in the number of women who participate in running events – mainly due to its ease of access combined with its minimal financial and social constraints^77^. Moreover, people may choose to participate in running events not only for the physical health benefits of such PA, but also for the good of their mental well-being and the positive socio-psychological effects^77^, as runners experience adrenaline, pleasure, relaxation, and more^78^.

Studies suggest that changes in the neural reward system are affected by the menstrual cycle^79^, and indeed, differences in the *COMT* rs4680 genotype prevalence might be involved in causing variable effects of estrogens^80^. As such, it could be hypothesized that the variable menopause status, which indicates female sex hormone levels, could be a significant interfering variable in the relationship between *COMT* rs4680 genetic polymorphism and PA frequency. In the current study we found that the differences in WEF and PA indexes according to *COMT* rs4680 polymorphism, as described above, exist only among women with regular period. Further research is needed in order to explore the relations between sex hormonal changes throughout the menstruation and PA frequency.

Despite the contribution of this study to the literature, this study has several limitations. First, the sample size was relatively small. Women are seriously under-investigated and future research could benefit from expanding on this exploratory study through larger sample sizes, including larger groups of sedentary participants. In addition, PA was measured through a self-reported questionnaire. Studies indicate an overestimation of PA when reporting is performed subjectively^26^. That being said, this bias was consistent throughout the sample, and as such decreases this limitation. Finally, dopamine levels were not measured, nor were female sex-hormones levels of reward, emotions, or motivation for engaging in PA. Thus, future studies could strongly benefit from focusing on the biological mechanism that underlies the relationship between the genetic polymorphism that is associated with the reward system and with PA engagement.

In conclusion, it seems that unlike women who are genetically predisposed towards neurological rewards from PA engagement, others might need external rewards and motivation in order to become and stay active.

AR: Active Runner
ANR: active non Runners
WEF: Weekly Exercise Frequency
OAI: Occupation Activity Index
SAI: Sport Activity Index
LAI: Leisure time Activity Index
RI: Running Index

## Footnotes

1 Recommended levels of physical activity for health: Adults aged 18–64 years = at least 150 minutes of moderate-intensity aerobic physical activity throughout the week, or at least 75 minutes of vigorous-intensity aerobic physical activity throughout the week (or an equivalent combination); aerobic activity should be performed in bouts of at least 10 minutes each session; for additional health benefits, adults should increase their moderate-intensity aerobic PA to 300 minutes per week or engage in 150 minutes of vigorous-intensity aerobic PA per week (or an equivalent combination); muscle strengthening activities should be performed using major muscle groups at least twice a week^81^

